# Solving the riddle of the evolution of Shine-Dalgarno based translation in chloroplasts

**DOI:** 10.1101/440842

**Authors:** Iddo Weiner, Noam Shahar, Pini Marcu, Iftach Yacoby, Tamir Tuller

**Affiliations:** School of Plant Sciences and Food Security, The George S. Wise Faculty of Life Sciences, Tel Aviv University, Ramat Aviv, Tel Aviv 69978, Israel; Department of Biomedical Engineering, The Iby and Aladar Fleischman Faculty of Engineering, Tel Aviv University, Tel Aviv 6997801, Israel; The Sagol School of Neuroscience, Tel Aviv University, Tel Aviv 6997801, Israel

**Keywords:** Shine-Dalgarno, Chloroplast, Ribosome-binding, Translation initiation

## Abstract

The chloroplast, a photosynthetic organelle found in all plant and algae species, originates from an ancient event in which a cyanobacterium was engulfed by a larger eukaryote. Thus, modern chloroplasts still harbor a bacterial-like genome and carry out all stages of gene expression, including mRNA translation by a 70S ribosome. However, the Shine-Dalgarno model, which predominantly regulates translation initiation by base-pairing between the ribosomal RNA and the mRNA in model bacteria genera, was reported to have ambiguous effects on chloroplast gene expression. Here we show that while the Shine-Dalgarno motif is clearly conserved in proteobacterial mRNAs, its general absence from chloroplast mRNAs is observed in cyanobacteria as well, promoting the idea that the evolutionary process of reducing the centrality of the Shine-Dalgarno mechanism began well before plastid endosymbiosis. As plastid ribosomal RNA anti-Shine-Dalgarno elements are highly similar to their bacterial counterparts, these sites alone cannot explain the decline in plastid Shine-Dalgarno generality. However, by computational simulation we show that upstream point mutations modulate the local structure of ribosomal RNA in chloroplasts, creating significantly tighter structures around the anti-Shine-Dalgarno locus, which in-turn reduce the probability of ribosome binding via the Shine-Dalgarno mechanism. To validate our model, we expressed a mCherry gene harboring a Shine-Dalgarno motif in the *Chlamydomonas reinhardtii* chloroplast. We show that co-expressing it with a 16S ribosomal RNA, modified according to our model, significantly enhances its expression compared to co-expression with an endogenous 16S gene.

**Significance statement:** Chloroplasts are fascinating intracellular organelles which have evolved from an ancient cyanobacterium engulfed by a larger eukaryote. Surprisingly, the canonical mechanism regulating bacterial translation initiation – Shine-Dalgarno – has been shown to play a reduced role in chloroplasts. Here, we show that mutations upstream from the anti-Shine-Dalgarno element decrease the probability of spontaneous ribosome binding by modulating the secondary structure of the ribosomal RNA. These mutations constitute a regulatory step which acclimates the Shine-Dalgarno mechanism to the translational regulation regime of chloroplasts. Interestingly, we show that these chloroplast features occur in modern cyanobacteria as well, promoting the idea that they have evolved prior to endosymbiosis.

## Introduction

Chloroplasts are the intracellular plant organelles in which photosynthesis takes place. The origin of these organelles is an early event of endosymbiosis in which an ancient cyanobacterium was engulfed by a eukaryotic host (1, 2). During the adaptation to this new symbiosis, the majority (∼95%) of plastid genes have been horizontally transferred to the nucleus. However, modern chloroplasts maintain a small circular genome consisting of ribosomal RNAs (rRNA), transfer RNAs, and open reading frames coding for ribosomal proteins, elongation factors, photosynthetic complexes and other housekeeping functions (3). On average, roughly 100 coding sequences (CDSs) are translated within chloroplasts by a bacterial-like 70S ribosome and a matching set of gene expression machinery (4, 5).

While translation initiation (TI) has been described as a major rate-limiting step for overall chloroplast gene expression (4, 6, 7), its dynamics and similarity to bacterial TI remain unclear. The canonical model for explaining TI in prokaryotes is the Shine-Dalgarno (SD) mechanism in which the 30S ribosomal subunit is recruited to the mRNA through base-pairing between a mRNA motif (SD sequence) found just upstream of the START site and an anti-SD (aSD) conserved sequence found at the 3’ edge of the 16S rRNA (8, 9). In this TI model, the proximity of the binding site to the START codon (roughly 10 base-pairs upstream) is a key feature necessary for proper initiation (10–13) (Fig. 1*A*).

**Fig. 1.**
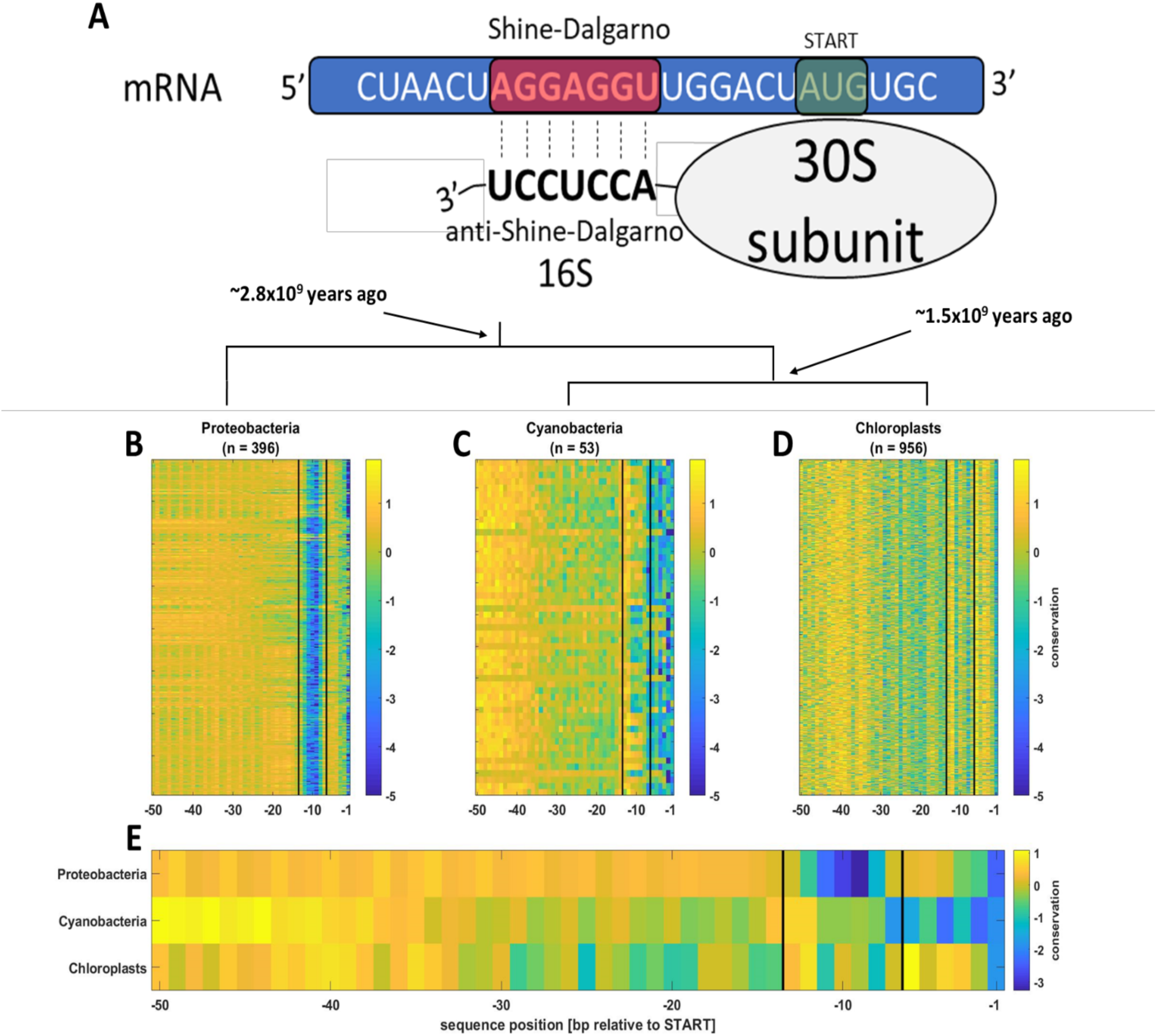
The Shine-Dalgarno motif does not show a clear signal in both chloroplast and cyanobacterial mRNAs. (*A*) Schematic illustration of the Shine-Dalgarno mechanism. A conserved element near the 3’ edge of the rRNA spontaneously binds a mRNA motif located slightly upstream of the START site by base-pairing. This interaction properly positions the small ribosomal subunit on the start codon to initiate translation. (*B,C,D*) Sequence conservation along the 5’ UTR. In each group the rows represent a single organism, selected to represent its genus. Lower values report on higher conservation in the specific position. The space between the vertical black lines represents the canonical location of the Shine-Dalgarno motif on the mRNA. The dendogram depicts the phylogenetic relations between the group. Node dating is based on phylogenetic reconstruction evaluations (2, 30, 31). (*E*) The mean value for each position across all rows in *B, C*, and *D*.

Since the availability of plastid sequences first began rising, the search for differences between chloroplast and bacterial TI features has attracted attention (4, 5, 7). While such differences often tend to be attributed to unique evolutionary adaptations taken on by chloroplasts post-endosymbiosis, computational works performed on several organisms have shown that the SD motif presence on mRNAs is decreased both in chloroplasts (14) and in cyanobacteria (10, 14–17). These data suggest that the decrease in SD centrality occurred prior to endosymbiosis. To present this on large-scale (Table S1), we calculated the 5’ UTR position-wise conservation for a representative from each sequenced chloroplast-containing genera, and compared the results to the equivalent cyanobacteria group of representatives. As an outgroup we used a similarly built database of proteobacteria – which are a large and diverse group, including key model organisms (*e.g. Escherichia coli, Helicobacter pylori, Pseudomonas aeruginosa, etc.*) from which the SD TI model was originally deduced. From this large scale analysis, it is evident that while proteobacteria show a clear SD signal centered roughly nine bases upstream from the START site (Fig. 1*B*), both chloroplasts and cyanobacteria show a noisier pattern with no clear signal in the SD area (Fig. 1*C,D*) (robustness testing against possible sequencing errors (18) is shown in Fig. S1). Since positioning plays a key role in the SD mechanism, this wide view observation provides robust evidence confirming that the SD mechanism in the ancient cyanobacterium engulfed during the formation of chloroplasts has already diverged significantly from its canonical role in proteobacteria. These data confirm that symbiosis with the nucleus (which applies to chloroplasts alone) could not be a main driver towards the reduced role of SD in chloroplasts. Alternatively, such drivers might include adaptation to oxygenic photosynthesis or to slower growth rates, as these features differentiate proteobacteria from cyanobacteria and chloroplasts (10, 19).

Previous experimental approaches aiming to clarify the functional role of the SD mechanism in chloroplasts yielded a mixture of results. While mutating a 5’ UTR SD motif is expected to reduce the gene’s translation efficiency, this outcome was only observed in several genes (20–22), whereas in others no significant effects were reported (21–24). In a recent report, Scharff *et al*. mutated the 16S aSD sequence and applied ribosomal profiling to quantify changes in translation rates across the tobacco chloroplast genome (25). Interestingly, it was found that SD binding is required for translation of only a subset of plastid genes, thus giving a broader context to previous reports.

However, it is well established that unlike in model bacteria, translation constitutes a major regulatory phase in chloroplast gene expression (5, 6, 26). This notion corresponds well with the decline in centrality of the spontaneous and energy-independent (4) SD TI mechanism (Fig. 1*E* and references above), which is more suitable for short-lived mRNAs (27–29) and regimes with strong transcriptional regulation. Thus, alterations in the plastid 16S rRNA 3’ edge that would add regulatory steps to the spontaneous SD mechanism and acclimate it to the chloroplast gene expression environment could be expected. In this work, we conduct a large scale analysis of these regions, from a systems-biology point of view, and find that mutation-based alterations in the ribosomal RNA structure affect SD-mediated TI.

## Results

As the 16S aSD sequence itself is highly conserved between chloroplasts and model bacteria (4, 5, 7), we hypothesized that structural alteration of the rRNA 3’ edge could constitute the additional regulatory step which acclimates the SD mechanism to the translation-regulation chloroplast regime. To test this hypothesis, we examined the 3’ edges of all 16S rRNAs in our database. While the aSD is clearly conserved in chloroplasts (and cyanobacteria), they contain a typical pattern of point mutations, clustered in two loci upstream of the aSD sequence (Fig. 2*A*). While these base substitutions do not directly affect the aSD sequence, they could change the local rRNA secondary structure, and thus indirectly complicate the aSD:SD binding event. To examine this hypothesis, we simulated the local folding of the 16S rRNA 3’ edge in all organisms in our database. To define the relevant region for simulation, we examined the 30S ribosomal subunit PDB structures from *Escherichia coli* (32) and the *Spinacia oleracea* chloroplast (33). We chose the region from nucleotide −28 relative to the aSD sequence until the 3’ edge, since unlike upstream regions which bind proteins and distant rRNA parts, in both organisms the RNA only interacts with itself in this area (Fig. 2*B*, Fig. S2). Interestingly, our folding simulations of this region show that the proteobacteria rRNA structures differ significantly from those of chloroplasts and cyanobacteria. To broadly quantify these effects across our database, we computed the rRNA folding that minimizes free energy for each organism and computed the number of aSD nucleotides found in structure. Our results clearly show that a significantly larger portion of aSD nucleotides is paired in chloroplasts and cyanobacteria (Fig. 2*C*). Since ribosome binding in the SD mechanism requires base pairing between the mRNA SD motif and the rRNA aSD sequence, the occupation of the latter by other bonds reduces its affinity to the mRNA and lowers the probability of binding; thus explaining the less canonical role that SD plays in chloroplast TI. Examining the most common structures received from each group reveals the loss of the well-described bacterial hairpin loop (34) in cyanobacteria and chloroplasts, and clearly emphasizes the structural differences in the aSD complex between groups (Fig. 2*D*). Four point mutations clustered in two loci (Fig. 2*A*, positions 4-5, 10-11) are the main drivers of this conformational change. Other point mutations and the slightly longer proteobacterial tails (Fig. 2*A*) have negligible effects on this observation (Fig. 2*D*). Interestingly, we also observed a significant positive correlation between the openness of the aSD element (*i.e.* the number of exposed aSD nucleotides) and the conservation of the mRNA SD motif within the proteobacterial group (Fig. S3). This observation indicates that the co-evolutionary dependence between these two traits occurs in proteobacteria as well.

**Fig. 2.**
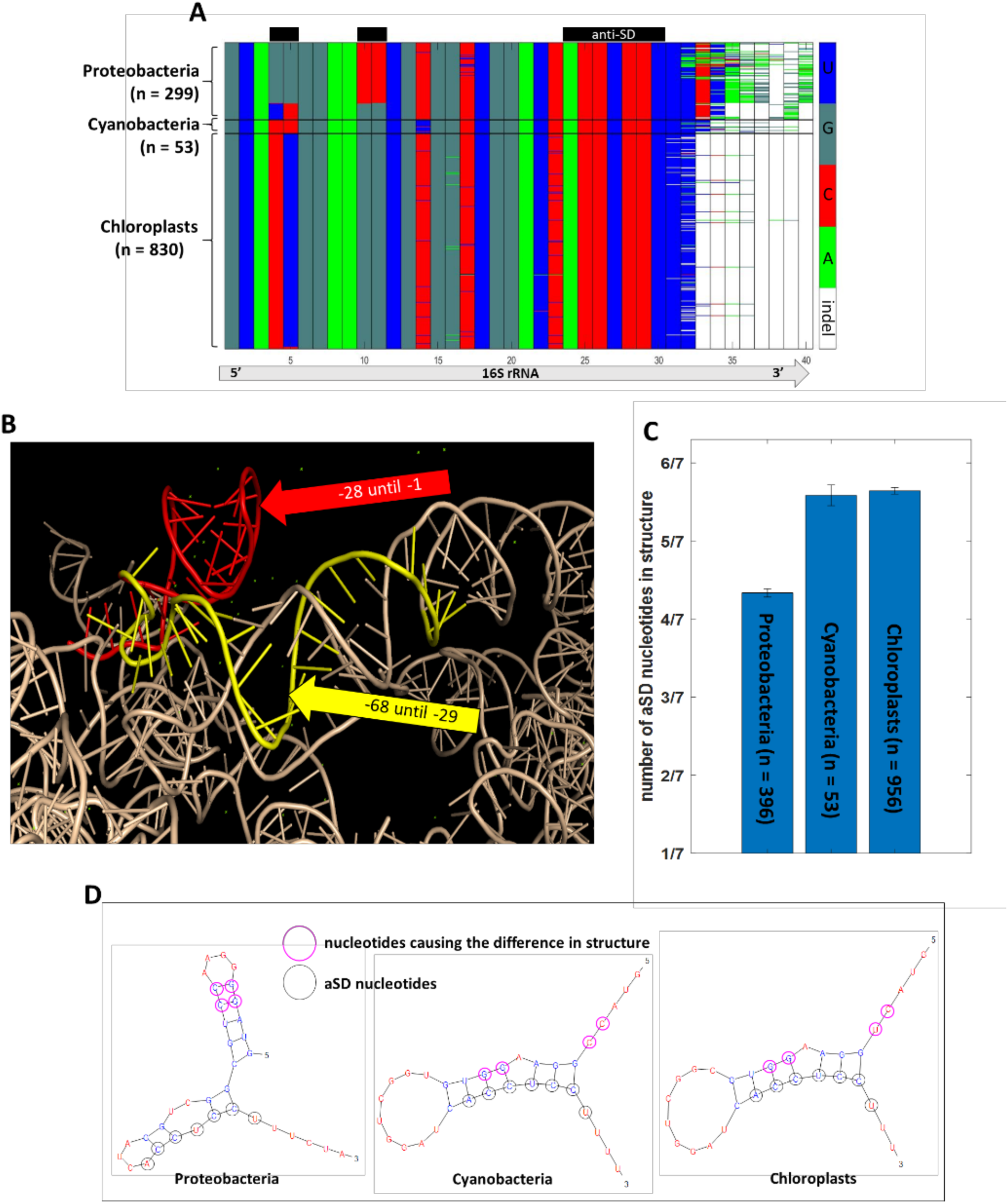
Local folding at the 16S 3’ ribosomal RNA edge affects the aSD structure. (*A*) Multiple sequence alignment of the 16S rRNA 3’ edge database, composed of a single representative species from each genus. The black boxes above the matrix highlight important regions: the anti-SD box represents the conserved region where this sequence is found, whereas empty boxes represent columns with high variability between the different groups. Several outlying species which introduce indels in the body of the alignment were excluded for visualization purposes. (*B*) Examination of the 3’ edge of the 16S rRNA in the 30S ribosomal subunit from the chloroplast of *S. oleracea*. The reference point is the anti-Shine-Dalgarno sequence (*e.g.* −1 means one nucleotide upstream from the aSD sequence). The 3’ edge of the molecule (red) only interacts with itself, while the region immediately upstream (yellow) interacts with distant parts of the rRNA. (*C*) After simulating the 16S 3’ edge secondary structure of each organism, the number of anti-SD nucleotides found in base-pairing interactions was computed. The bars show the mean ± SE of this value across all organisms in each of the groups. (*D*) Typical (most abundant) rRNA secondary structures. Anti-SD nucleotides are marked in dark green whereas the nucleotides that cause the structure discrepancy between groups are circled in pink.

According to this model, simply changing these two clusters of altered nucleotides into their proteobacterial form will modify the local folding at the 3’ edge of the 16S rRNA, and expose the aSD sequence to facilitate easier binding to SD motifs. To test this theory, we designed two vectors for chloroplast expression in the model green alga *Chlamydomonas reinhardtii*. Both coded for a codon optimized mCherry reporter protein with a clear SD motif properly situated upstream from the START codon. Vector A also coded the natural *C. reinhardtii* 16S rRNA gene, whereas vector B coded for a slightly modified version of this 16S gene in which the 3’ edge was mutated to match the canonical proteobacterial sequence (Fig. 3*A*). We transformed each of these vectors into the *C. reinhardtii* chloroplast, validated their proper integration by PCR, and brought all clones to homoplasmicity (Fig. S4, see *Methods* in *supplementary material*). Additionally, to confirm that the reporter gene mRNA levels were similar in all clones, we quantified mCherry transcript abundance by quantitative PCR (qPCR) (Fig. S5). Subsequently, we confirmed the presence of mCherry protein by immunoblotting (Fig. S6).

**Fig. 3.**
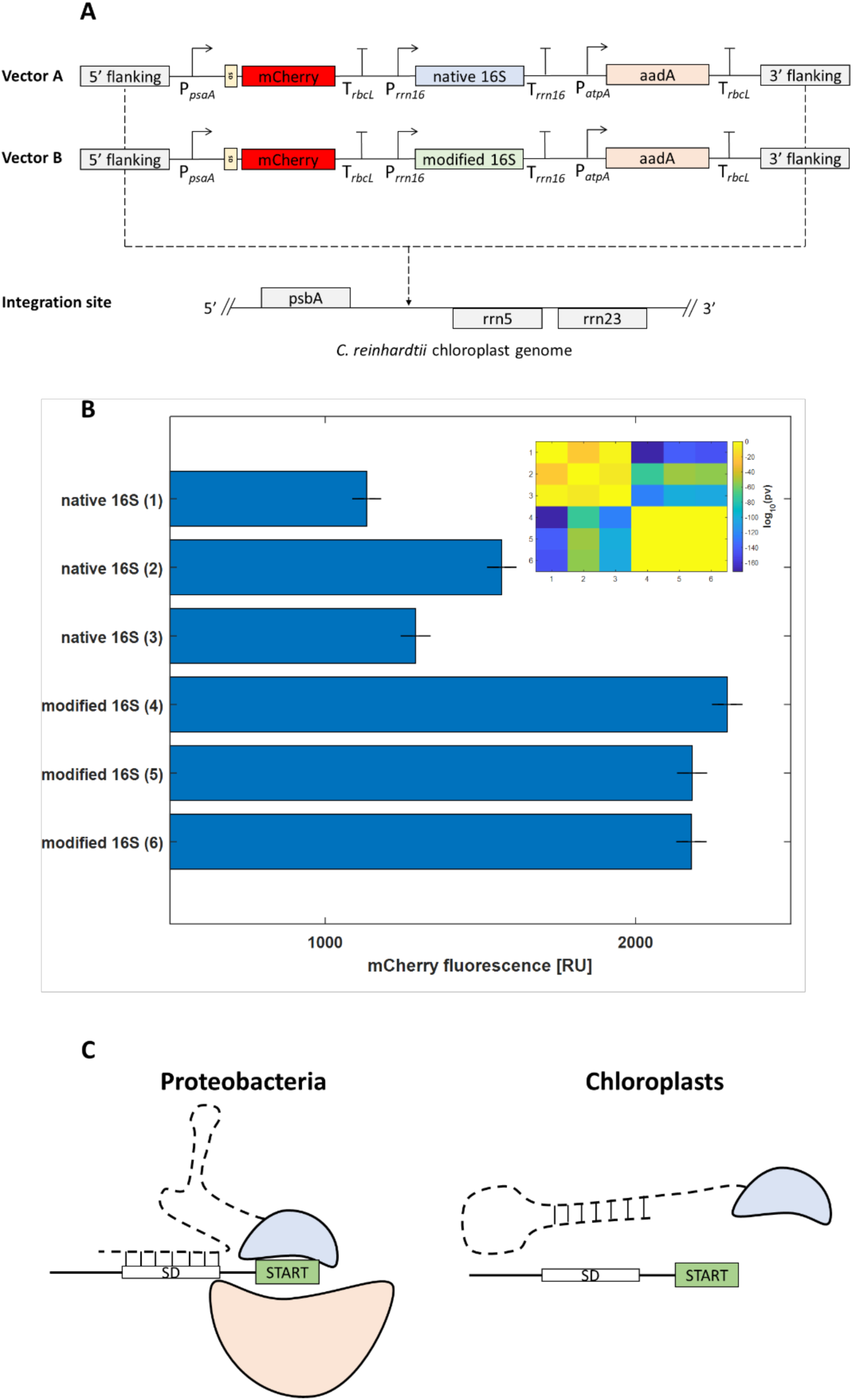
Introducing a modified 16S rRNA into the chloroplast genome enhances the expression of a reporter gene with a SD motif. (*A*) Schematic diagram of the chloroplast expression vectors. Vectors A and B were completely identical, except for point mutations at the 3’ edge of the 16S gene in vector B, which altered its secondary structure. (*B*) Mean mCherry fluorescence across 104 cells in three individual clones from each transformation group, as measured by flow cytometry. Data shown is mean ± SE. P-values in logarithmic scale are shown on the upper right. All p-values were computed using the two-tailed Wilcoxon rank-sum test. (*C*) Schematic illustration of the model representing the altered role of Shine-Dalgarno in chloroplast translation initiation; while bacterial 16S rRNAs are open and ready to base-pair with the mRNA, mutations in the chloroplast 16S 3’ edges tighten the structure of the aSD sequence and hamper spontaneous interactions with the mRNA.

To quantify the differences in translation efficiency between the two groups of transformants, we measured mCherry fluorescence for each of these clones by flow cytometry, in which 10^4^ single cells from each mid-log phase culture were examined. We observed that our reporter gene was significantly more highly expressed in the clones transformed with vector B (Fig. 3*B*). Since homologous recombination (HR) between the untouched endogenous 16S and the inserted synthetic 16S could potentially erase our modifications, we confirmed the presence of our point mutations on the synthetic 16S following the flow cytometry analysis (Fig. S7 [including further discussion on HR]). Alongside our computational analysis, this discrepancy in protein expression indicates that loose aSD structures are important for spontaneous aSD:SD base-pairing (Fig. 3*C*).

## Discussion

In this work we have shown that in cyanobacteria and chloroplasts, mutations upstream from the aSD element modulate the secondary structure of the rRNA 3’ edge, thus decreasing the probability for spontaneous ribosome binding through the SD mechanism. Recently, it has been shown that SD-mediated ribosome binding still occurs in the tobacco chloroplast, thus confirming that all machinery required for proper SD-mediated ribosome binding remained intact in plastids and still plays a role in controlling chloroplast TI (25). This explains the necessity of the aSD element, and together with our model it might also explain the conservation of the upstream mutations (Fig. 2*A*, positions 4-5, 10-11); according to this hypothesis, these mutations serve as part of an additional regulatory element which lowers the spontaneity of the aSD:SD interaction and acclimates the SD mechanism to the chloroplast environment in which regulation at the translational level is predominant. However, unlike in proteobacteria where SD is the canonical TI model, Scharff *et al*. have shown that it is a dominant TI mechanism for only a specific subset of genes. Importantly, this work shows that genes in which SD is essential for TI also tend to have strong mRNA secondary structures in the vicinity of their START sites. Together with our data, these observations could suggest that mRNAs controlled by SD might require a certain 5’ UTR secondary structure to unfold the 16S edge and reveal the aSD sequence in order to facilitate ribosome binding.

A useful by-product of this work is the establishment of a generalist method for enhancing heterologous expression in chloroplasts. Our baseline plasmid in this work, pLM21, is known to drive high chloroplast expression in *C. reinhardtii* (35–37), yet by adding a slightly modified copy of the 16S rRNA (Fig. 3*A*), we were able to roughly double the amount of protein achieved (Fig. 3*B*). As the aSD sequence is concealed in nearly all chloroplasts (Fig. 2), we expect that adding a modified 16S rRNA to any plastid expression vector would similarly enhance the translation of a target transcript harboring a SD motif in any chloroplast. However, one must take into consideration that: (i) while potential HR with the native 16S has not deleted the modified 16S version in any of our clones (Fig. S7), it could theoretically reverse the effect of the synthetic 16S over time, and (ii) while under our experimental conditions we observed no phenotypic difference between the transformation groups (Fig. S8), insertion of a mutated 16S rRNA could theoretically cause such differences in other organisms or under different conditions.

## Materials and Methods

*Further detail can be found in the SI Methods.

*We have placed several documents, containing various types of information, in the following depository:

https://github.com/iddoweiner/Solving-the-riddle-of-the-evolution-of-Shine-Dalgarno-based-translation-in-chloroplasts

Henceforth referred to this as *the depository*.

### Data Base Construction

To build the database used for the analyses described in this work, we retrieved the accessions for all organisms in NCBI that are classified as either proteobacteria, cyanobacteria or chloroplasts. The full list of accessions can be found in *the depository* (defined above). Within each group, we randomly selected one representative for each genus to appear in the final database. In Fig. 2*A* alone, several organisms were removed from the alignment as they introduced sporadic indels into the alignment matrix and interfered with data visualization.

### Conservation (Entropy Z-score)

To compute the organism-specific position conservation scores presented in Fig. 1 we initially obtained the 50 base-pairs (bp) upstream from the start site in all the CDSs (Fig. S9) and computed the position specific scoring matrix. For each matrix column we computed the Shannon Entropy by:

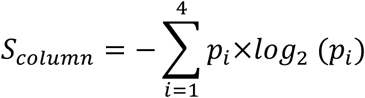

More specifically, given the frequencies p(A), p(C), p(G), p(U), the entropy S of a column (position) is:

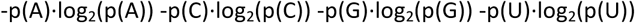

We computed the same entropy score for 100 different random models of the same matrix, in which the columns were permutated together to maintain the alignment. The final value shown in Fig. 1 is the margin between the entropy computed on the native 5’ UTRs and the average of those computed for the randomized UTRs, divided by the standard deviation of the random model distribution (Z-score).

### 16S rRNA local folding simulation

To simulate the local folding at the 3’ edge of 16S rRNAs, we first examined the 30S ribosomal subunit PDB structures from *Escherichia coli* (32) and the *Spinacia oleracea* chloroplast (33). While using the conserved aSD sequence as a reference point, we observed that from roughly nucleotide −28 until the 3’ edge, base pairing is scarce and importantly – only interacts within itself (Fig. 2*B*, Fig. S2). Regions slightly upstream from this point interact with distant self-regions (*e.g.* with regions located 500 bp upstream on the 16S), ribosomal proteins and other rRNAs. Thus, we selected this area for our simulations. The folding structure was computed using the ViennaRNA package (38).

### Shine–Dalgarno consensus sequence

While there is some debate whether the exact consensus SD sequence is AGGAGGU or just AGGAGG (thus the aSD is either ACCUCCU or CCUCCU, respectively), previous studies lean towards the former (39, 40). The high level of conservation of adenosine in position 24 (Fig. 2*A*) could strengthen this notion. While we have indeed used the 7-base consensus sequence in this work, the only result that depends upon the exact consensus sequence is the one presented in Fig. 2*B*. A similar analysis based upon the 6-base consensus sequence is shown in (Fig. S10), where the conclusion remains the same.

### Algal cultures and chloroplast transformation

*Chlamydomonas reinhardtii* wild-type (D66) strains were grown in Tris-Acetate-Phosphate medium at 25°C under continuous cool daylight and cool white fluorescent lights (90 µE m^-2^ s^-1^) stirring in 100 mL Erlenmeyers capped with silicone sponge enclosures, as previously described (41, 42). Vectors A and B (Fig. 3*A*) were cloned into the pLM21 (35) and transformed into the *C. reinhardtii* wild-type chloroplast using a biolistic gun. The full annotated plasmid maps appear in *the depository* (defined above). Several rounds of PCR were used to isolate homoplasmic clones (36, 43) (also see Supplementary Methods).

### Flow Cytometery

The mCherry fluorescence was quantified from a 200 µL sample in mid-log phase (10^6^ cells mL^-1^) from each positive clone, using the S1000EXI Benchtop Flow Cytometer (https://stratedigm.com/s1000exi-flow-cytometer). The mCherry parameters were selected according to the manufacturer’s instructions. The auto-fluorescence of the *C. reinhardtii* wild-type was measured in the same experiment and subtracted from the clones’ intensity.

### Code availability

The code generated during the current study is available from the corresponding author on reasonable request.

## Acknowledgements

This study was supported in part by: The Israeli Ministry of Science Technology and Space; Israel Science foundation 1646/16; NSF-BSF Energy for Sustainability 2016666; A fellowship from the Edmond J. Safra Center for Bioinformatics at Tel-Aviv University; A fellowship from the Manna center for Plant Biosciences; A fellowship from the Rieger foundation for Environmental studies. We would like to thank: The G. Peltier lab for kindly sharing their pLM21 vector; Mrs. Barel Weiner for critical reading; Mrs. Orit Sagi-Assif for Flow Cytometry support; The Adi Avni lab for providing us purified mCherry. IW, NS, IY and TT designed the research; IW and NS performed the experimental and computational procedures; PM analyzed the PDB ribosome structures; IW, NS, IY and TT wrote the paper. All authors declare no conflict of interest. No conflicts, informed consent, human or animal rights applicable. The datasets generated during and/or analyzed during the current study are available from the corresponding author on reasonable request.

## Supplemental information

<u>*Important note:</u>

We have placed several documents, containing various types of information, in the following depository:

Henceforth referred to this as ***the depository***.

### Supplemental Methods

#### DNA extraction

Total DNA was extracted from 50 mL mid-log phase *C. reinhardtii* (3 × 10^6^ cells mL^-1^) cultures. Cultures were centrifuged (2800g for 7 min) and 30 mg of the cell pellet was taken for total DNA extraction using an OMEGA E.Z.N.A plant DNA kit.

#### Quantitative PCR

Total RNA was extracted from 50 mL mid-log phase *C. reinhardtii* (3 × 10^6^ cells mL^-1^) cultures. Cultures were centrifuged (2800g for 7 min) and 100 mg of the cell pellet was taken for total RNA extraction using RNeasy plant Mini Kit (QIAGEN 74903). The cell suspension was lysed in a Minilys tissue lyser for 90 seconds (Bertin technologies). Total RNA was treated with Turbo DNA-free™ Kit (Ambion AM1907) according to the manufacturer’s recommendation, followed by PCR and gel electrophoresis quality check. The RNA concentration was determined using a NanoDrop® ND-1000 Spectrophotometer. 2 µg of purified RNA from each sample were used for complementary DNA (cDNA) synthesis using High capacity cDNA Reverse Transcription Kit (ABI 4368814) that was performed with random primers according to the manufacturer protocol.

The qPCR reaction was performed with Applied Biosystems StepOnePlus™ real time PCR system and software using Fast SYBR Green Master Mix (ABI 4385612) as the fluorescence reporter. Each reaction had a 10 μl final volume containing: 2µl DNA template (plasmid DNA dilutions for the standard curve or 30 ng cDNA for measuring a clone’s transcript abundance), 200nM primers (forward and reverse) and 5µl SYBR Green Master Mix. Reaction conditions included 40 amplification cycles according to ABI StepOnePlus™ system and software default; 3 s at 95 °C and 30 s at 60 °C. Each reaction was performed in technical triplicates and the average was considered as one measurement. Primers for mCherry and the reference gene were designed using an in-home algorithm based upon the Primer-BLAST designing tool (NCBI) to eliminate the option for gDNA amplification. Ct was calculated under default settings for the real-time sequence detection software (StepOne™ Software v2.3, Applied Biosystems). A sample from all cDNA pools (each isolated from a different clone) were put through a parallel qPCR analysis, amplifying a sequence of the reference gene *cblP* (44).

#### Equations

1. ΔC_T_ = C_T_ (*mCherry*) - C_T_ (*cblp*)
2. Normalized transcript abundance = 2-ΔC_T_
3. ΔΔC_T_ = ΔC_T_(vectorA) - ΔC_T_(vectorB)
4. Transcript fold change = 2-ΔΔC_T_

#### Immunoblot assay

Soluble proteins were isolated from 50 mL mid-log phase *C. reinhardtii* (3 × 10^6^ cells mL^-1^) cultures. A volume that corresponded to 140 µg chlorophyll was concentrated (2800g for 7 min) and taken for induction. Immediately thereafter cells were precipitated (3200×*g*, 5 min) and re-suspended in buffer A (50 mM Tris–HCl pH 8.5, 20 mM Na-dithionite, 60 mM NaCl and 1 mM protease inhibitor cocktail). The cell suspension was lysed in a Minilys tissue lyser (Bertin technologies) by three 5,000 rpm cycles of 45 seconds each in the presence of Sigma glass beads (425–600 µm). The soluble proteins were isolated by centrifugation (10 min, 4°C, 14,000×*g*) and the total soluble protein concentration was determined by commercial Bradford solution (Bio-Rad). The protein concentrations were all in the range of 1.84 ± 0.26 µg µL^-1^. After adding sample buffer (Bolt™ LDS Sample Buffer) and reducing agent, samples were boiled at 95°C for 4 min. 20 µg of soluble proteins were loaded on 4–12 % Bis–Tris Plus PAGE gels (Novex by Life technologies) and analyzed by immunoblotting using iBind™ blotter and its specific blocking reagents (Life technologies). As a primary antibody we used a rabbit polyclonal mCherry antibody (https://www.biovision.com/mcherry-antibody.html). Membrane images were taken using a DNR-MicroChemi station whereas the size markers were overlaid using the RGB camera included in the DNR. Ponceau staining was applied immediately after the immunoblot assay, and the stained membrane image was taken using a UVITEC Cambridge gel imaging system.

#### Primers

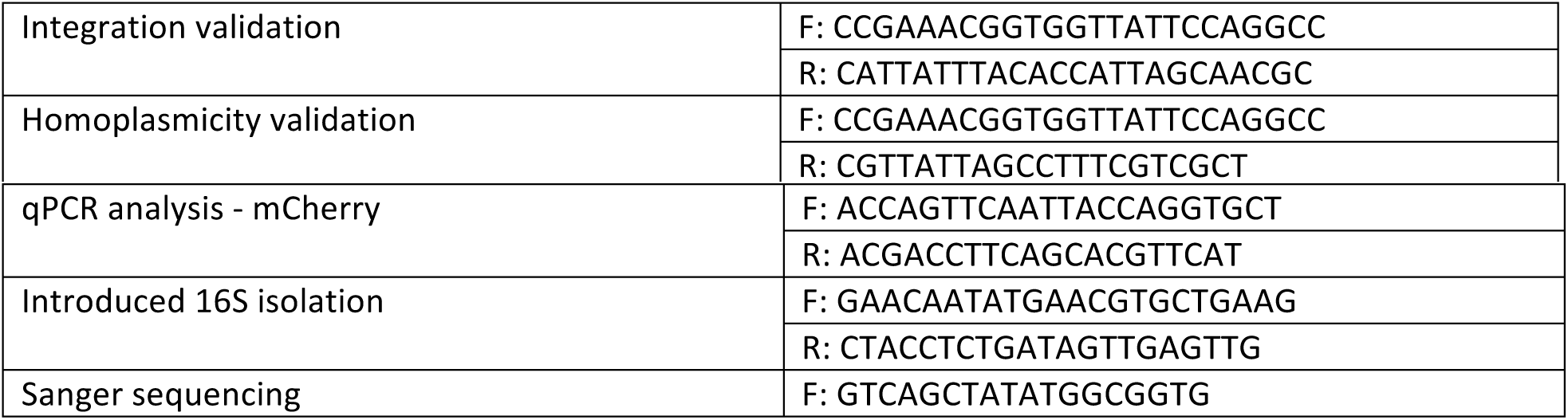

**Table S1:** Comparison between papers that studied the global signature of SD on cyanobacteria and chloroplast mRNAs.

| <u>Paper title</u> | <u>#Cyano organisms</u> | <u>#chloroplast organisms</u> | <u>Total number of organisms</u> |
| --- | --- | --- | --- |
| Correlations between Shine-Dalgarno Sequences and Gene Features Such as Predicted Expression Levels and Operon Structures | 1 | 0 | 30 |
| Local Absence of Secondary Structure Permits Translation of mRNAs that Lack Ribosome-Binding Sites | (6,200 sequences) | (11,238 Sequences) | (63,595 Sequences) |
| Predicting Shine–Dalgarno Sequence Locations Exposes Genome Annotation Errors | 2 | 0 | 18 |
| Comparative transcriptomics of two environmentally relevant cyanobacteria reveals unexpected transcriptome diversity | 1<br>(2 different strains) | 0 | 1 |
| Dynamic evolution of translation initiation mechanisms in prokaryotes | 9 | 0 | 277 |
| **Our current manuscript** | **53**<br>**(265,989 Sequences)** | **956**<br>**(83,513 Sequences)** | **1,405**<br>**(1,952,270 Sequences)** |

**Fig. S1:**
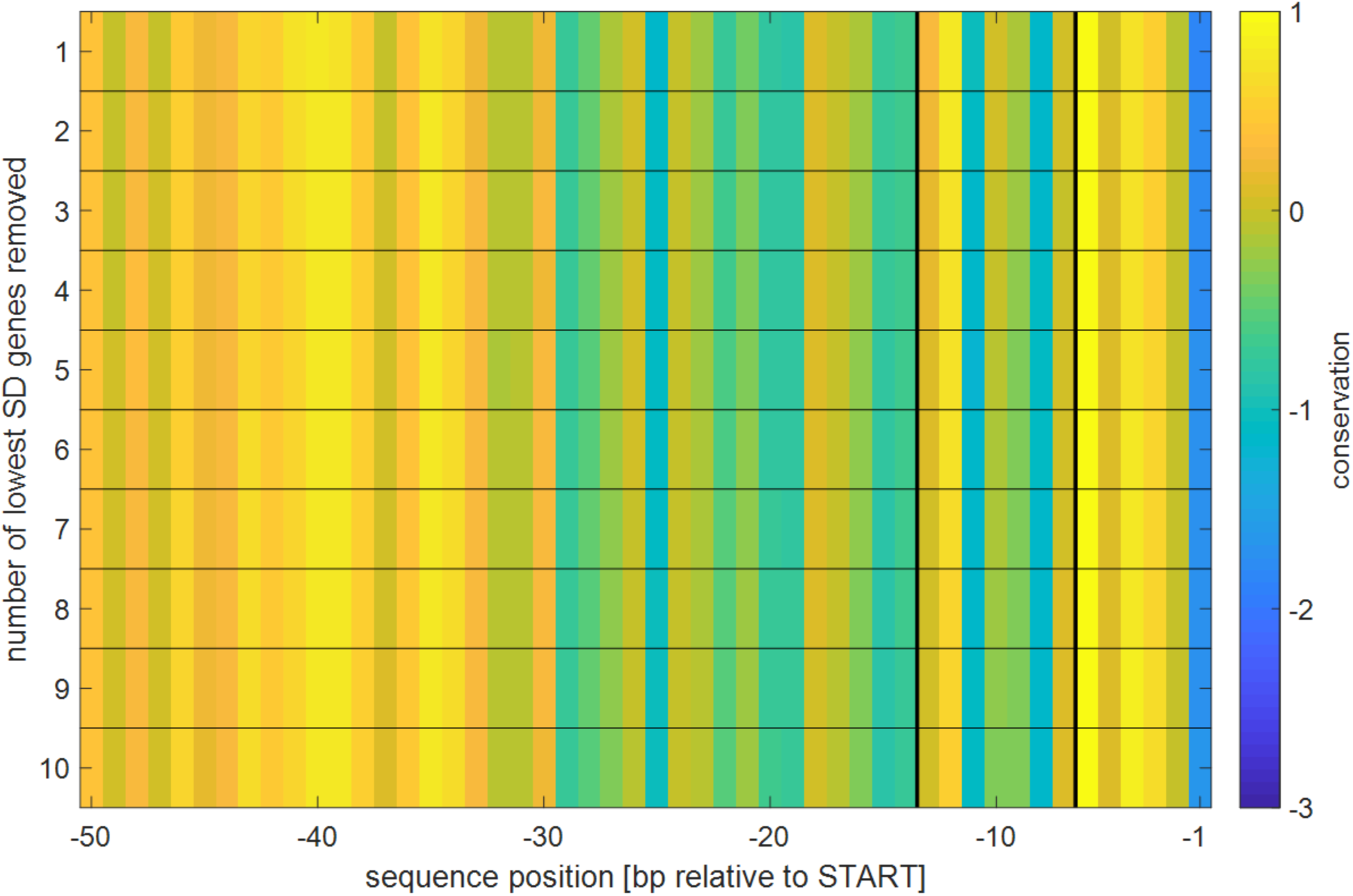
Robustness test against plastome sequencing/annotation errors. In this analysis we examined the robustness of the observation in Fig. 1*E*, given possible sequencing / annotation errors in plastomes, such as those described in Gallaher *et al* (18). To validate that the absence of the SD motif is not an artifact caused by such errors (which could introduce false START codons into the genomic record files and raise an alternative explanation to the absence of SD signal in chloroplasts), we conducted an iterative robustness exclusion process: in each iteration we omitted the 5’ UTR with the lowest SD score (*i.e.* the one that best supports the observation in Fig. 1*E*) from each plastome in our database, and then re-ran the conservation calculations. At the end we averaged these values across all organisms and drew a new heat-map in a similar manner to the heat-map presented in Fig. 1*E*. We iterated this process until 10 genes were removed from each plastome - simulating a case where in each plastome the 10 mRNAs with the lowest SD score are a result of a sequencing / annotation errors. The similarity between these results and the those shown in Fig. 1*E* emphasize the robustness of this result. The numeric values matrix can be found in ***the depository*** (‘Position-specific conservation values received from omission experiment.xlsx’).

**Fig. S2:**
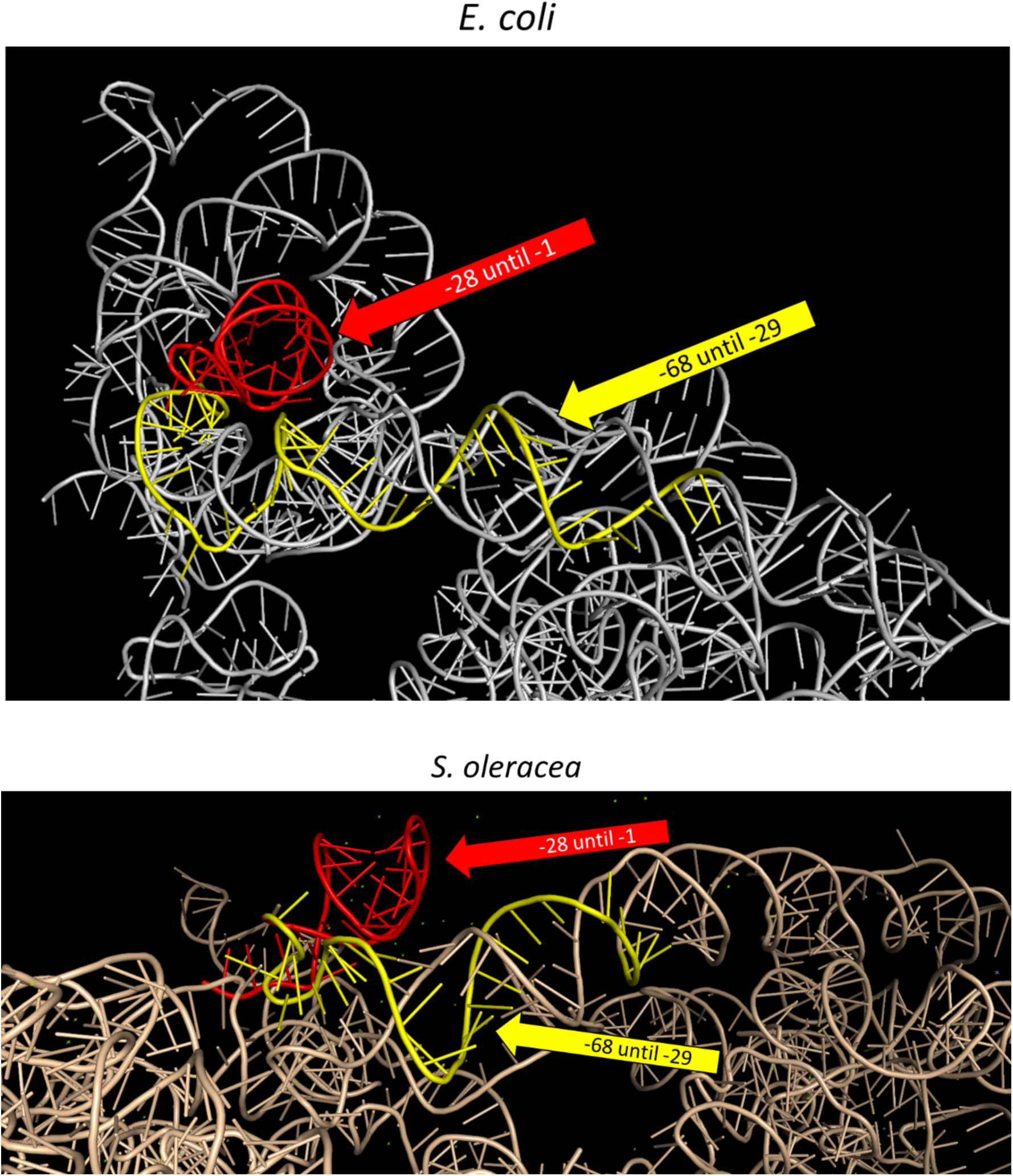
Analysis of the 16S 3’ edge 3D structure. In this analysis we examined the 3’ edge of the 16S rRNA in the 30S ribosomal subunit from *E. coli* (32) and from the chloroplast of *S. oleracea* (33) (no high resolution structure of a cyanobacterial 30S ribosomal subunit could be found on PDB at this time). We used the highly conserved aSD as a reference point (*e.g.* −1 means one nucleotide upstream from the aSD sequence). In both organisms while the 3’ edge of the molecule (red) only interacts with itself, the region immediately upstream (yellow) interacts with distant parts of the rRNA and with proteins (not shown here for clearer visualization). The PDB validation reports can be found at ***the depository***.

**Fig. S3:**
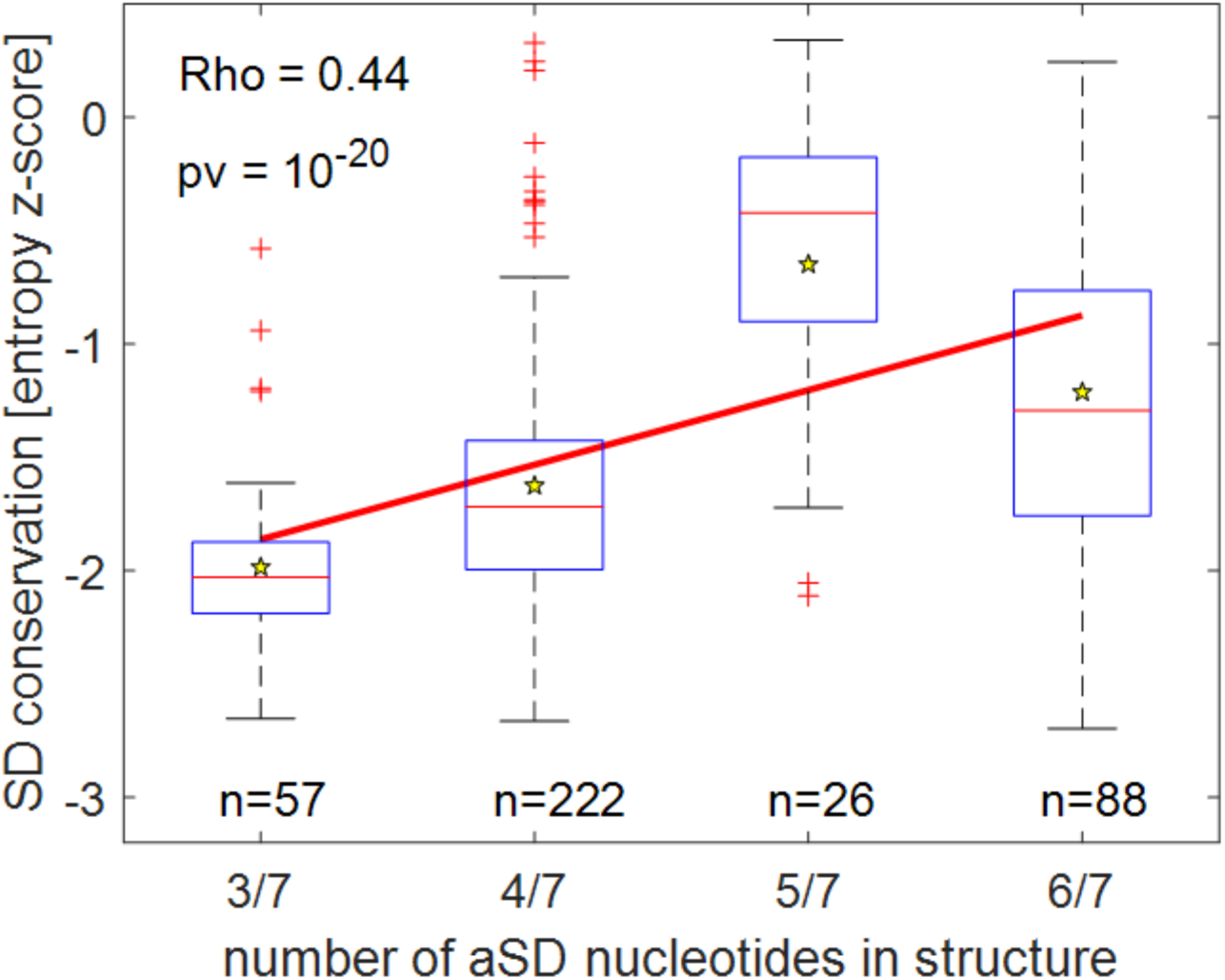
Correlation between the conservation of mRNA Shine-Dalgarno motifs and the structure of the 16S rRNA in proteobacteria. This analysis focuses on the proteobacterial group alone: we simulated the secondary structure of the 16S 3’ edge and computed the number of aSD nucleotides found in base-pairing interactions; these data are shown on the X-axis. For each of these organisms, we also computed the average position-specific conservation within the mRNA SD area (*i.e.* for each organism - the average value from inside the boundaries shown in Fig. 1*B*); these data are shown on the Y-axis. It is important to note that since these values are given in entropy Z-scores, lower values signify higher conservation. The boxplots show the median conservation and its distribution, whereas the yellow star signifies the average. The number of organisms (n) in each discrete X-axis value is given. Three outlying species (with 1/7 and 7/7 aSD nucleotides in structure) were removed from this graph to facilitate visualization, but they were included in the calculation of the correlation coefficient (Spearman’s correlation coefficient: 0.44, p=10^-20^), and in the calculation of the least squares line, shown in red. This analysis shows that open aSD structures are significantly correlated with conserved SD mRNA motifs, as could be expected from the model illustrated in Fig. 3*C*.

**Fig. S4:**
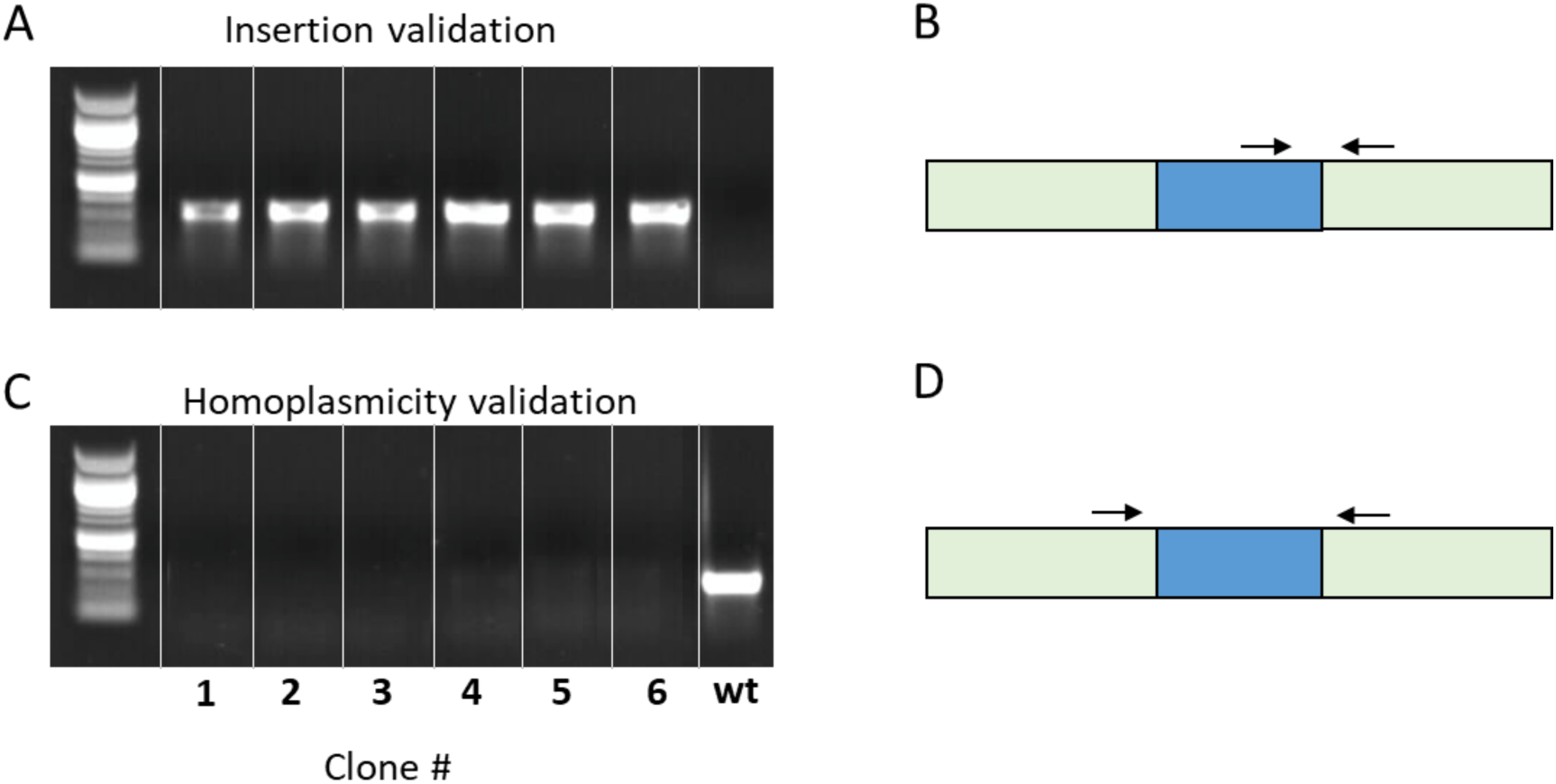
Conformation of gene integration and isolation of homoplasmic strains. PCR using whole cell lysates as described in Rasala *et al* (43). In (*A*) the vector integration was validated by PCR, where the forward primer annealed to the insert and the reverse primer annealed to the plastome, adjacent to the planned integration site, as depicted in (*B*). In (*C*) homoplasmicity was tested by PCR, where each primer flanked the integration site, as depicted in (*D*). Given the substantial length of the insert and the short synthesis time used in the reaction, bands are only received for un-engineered plastome copies. The lack of bands indicates homoplasmicity. All bands presented here were obtained in the same reaction, which had 40 PCR cycles.

**Fig. S5:**
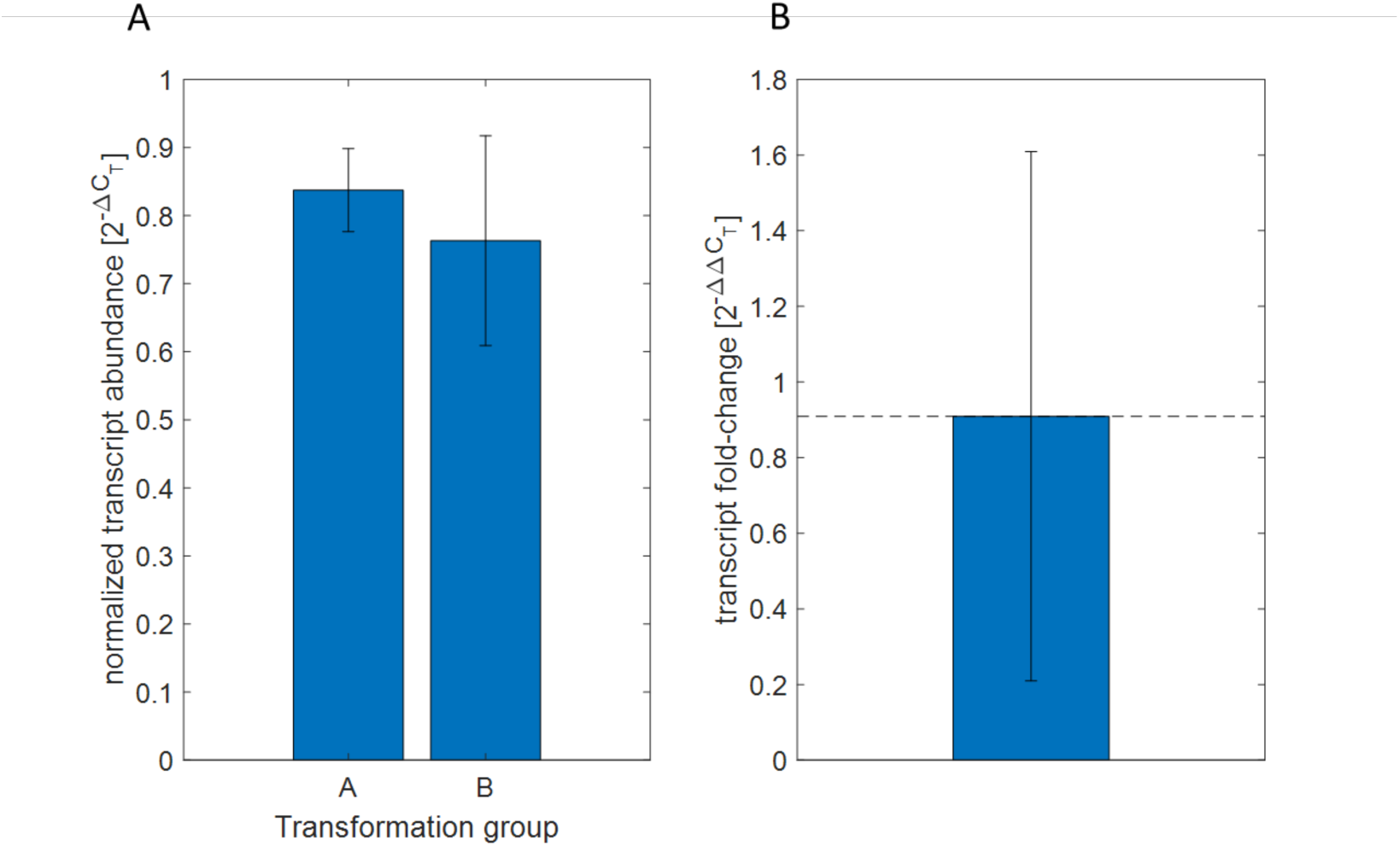
Relative quantification of mCherry transcripts. In this analysis we quantified the abundance of mCherry mRNA in all six clones. To this end, we extracted total RNA and reverse-transcribed it to create a cDNA library. Both mCherry and a reference gene (*cblP* (41, 44)) were amplified in a quantitative PCR reaction, and the ΔC_T_ values were calculated for each clone. Panel (*A*) shows the average ΔC_T_ values for clones transformed with vectors A and B; no significant difference between these groups was found (p = 0.68 [ttest], p = 0.7 [Wilcoxon’s rank-sum]) Data shown is mean ± SE. Panel (*B*) depicts the ΔΔC_T_ value, which quantifies the fold change in expression between the groups. The error bars given here are the 95% confidence interval. See Supplemental Methods for a detailed explanation of the procedures.

**Fig. S6:**
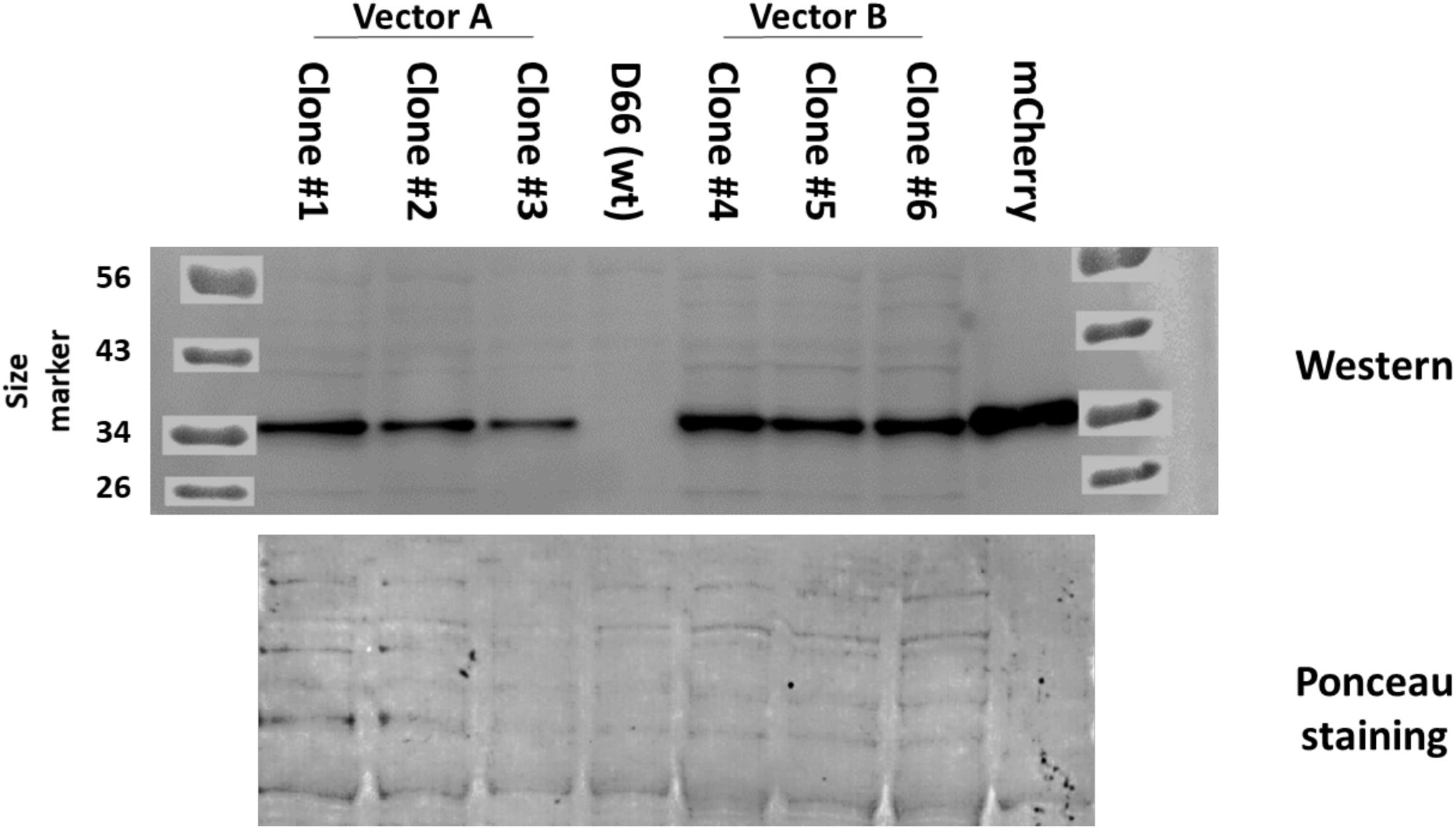
Confirmation of recombinant mCherry presence by immunoblotting. In this analysis we confirmed the presence of the inserted mCherry in all our clones, at the protein level. To this end we isolated the total soluble protein content from each clone, and loaded equal amounts in each lane (calculated according to chlorophyll, see Supplemental Methods). Our primary antibody detected mCherry in all lanes, except in the wild-type (upper panel). The positive control in the right lane is mCherry extracted from *Solanum lycopersicum* following tDNA insertion. Ponceau staining was performed on the membrane after the luminescence exposure to visualize the loading quantities. See Supplemental Methods for a detailed explanation of the procedures.

**Fig. S7:**
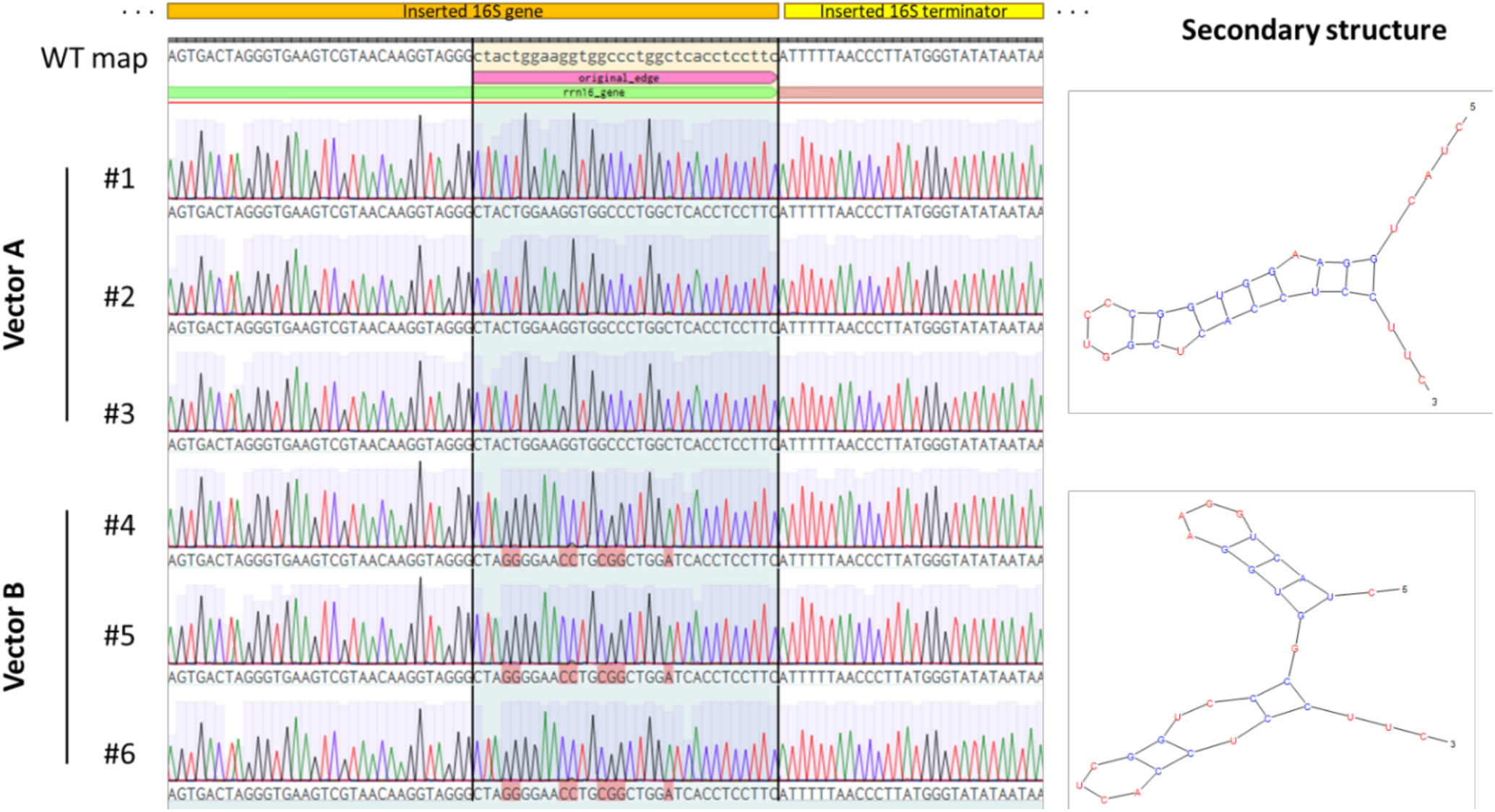
Targeted sequencing of the point mutations introduced to the synthetic 16S rRNA 3’ edge. To ascertain the presence of our modifications on the inserted 16S copy, we extracted DNA from all strains (1-6) and isolated the region which is supposed to contain the 3’ edge of our inserted 16S gene. We used Sanger sequencing to analyze this region and aligned it with the *C. reinhardtii* plastome reference sequence we used. Thus, we confirmed the presence of our mutations in all clones transformed with vector B (clones #4, #5, #6). Notably, as the *C. reinhardtii* chloroplast contains roughly 83 plastome copies, reversion of a small portion of copies within a cell culture to their WT form could theoretically happen. This analysis, taken together with the data shown in Fig. 3, proves that no significant reversion has occurred.

**Fig. S8:**
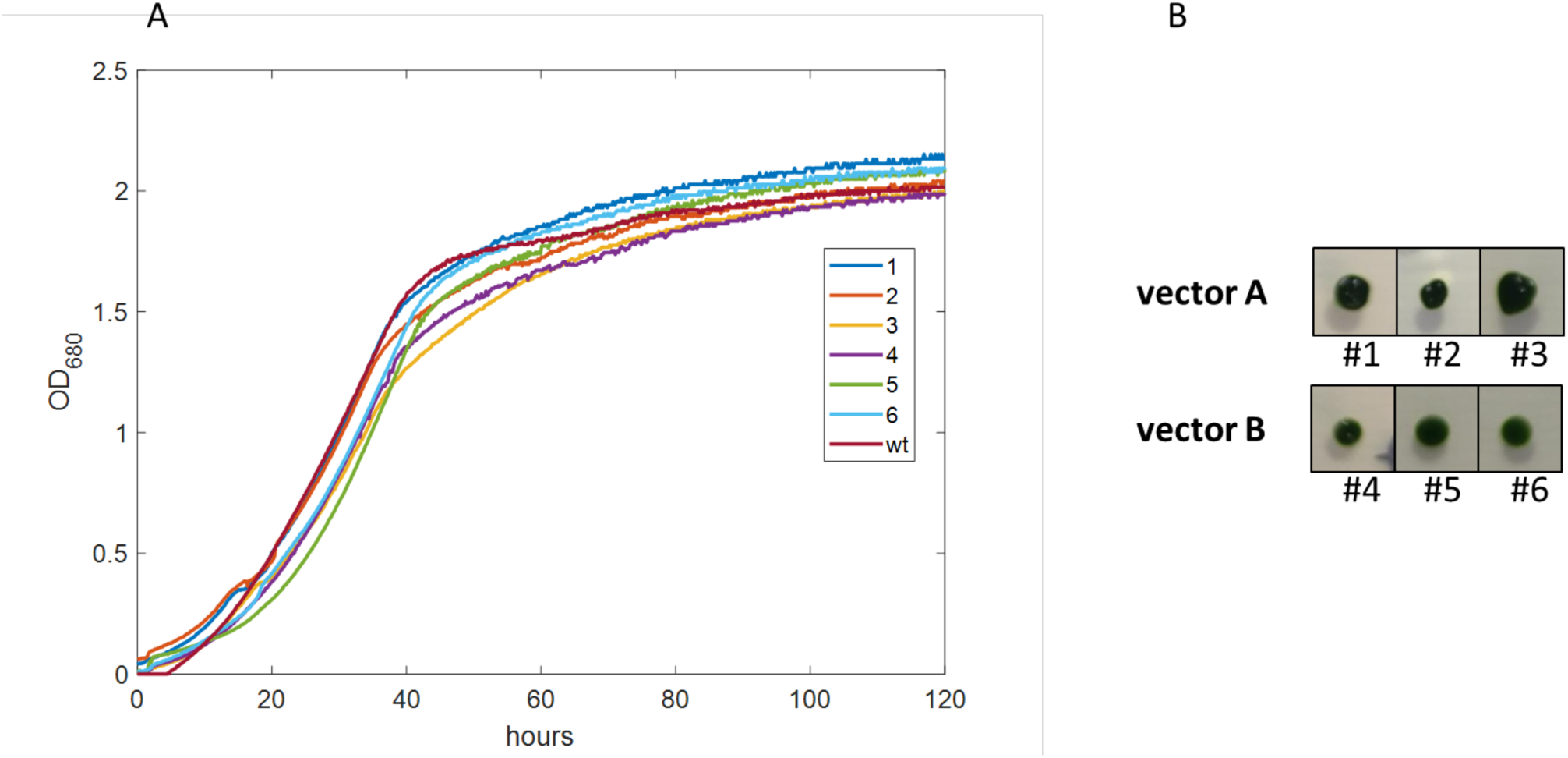
Phenotypic similarity between clones from different transformation groups. (*A*) Cultures in mid-log phase were diluted to 0.1 OD_680_, and cultured in a Multicultivator (MC 1000 / OD, PSI) with constant bubbling. Irradiance was set to 190 µE and OD_680_ was measured in 10 minute intervals. (*B*) Colonies growing on Tris-Acetate-Phosphate medium containing spectinomycin as a selection marker, at 25°C under continuous cool daylight and cool white fluorescent lights (90 µE m^-2^ s^-1^).

**Fig. S9:**
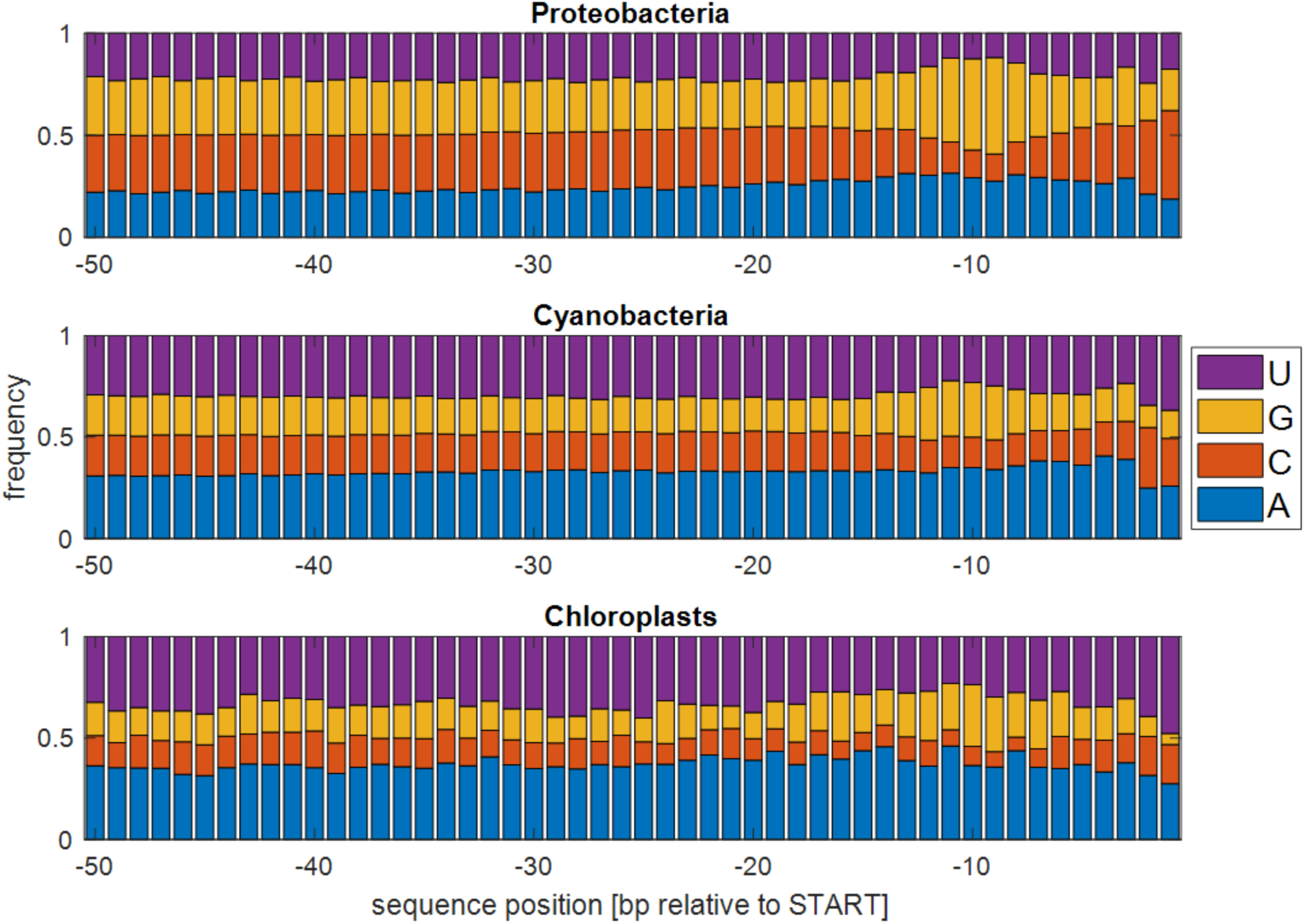
Nucleotide frequencies in 5’ UTRs. The plain PSSM for all organisms in our database is shown here. As opposed to Fig. 1, where the value given is compared to a random model which eradicates noise caused by high/low GC content, here these biases are conspicuous. n = 396 (Proteobacteria), 53 (Cyanobacteria), 956 (Chloroplasts).

**Fig. S10:**
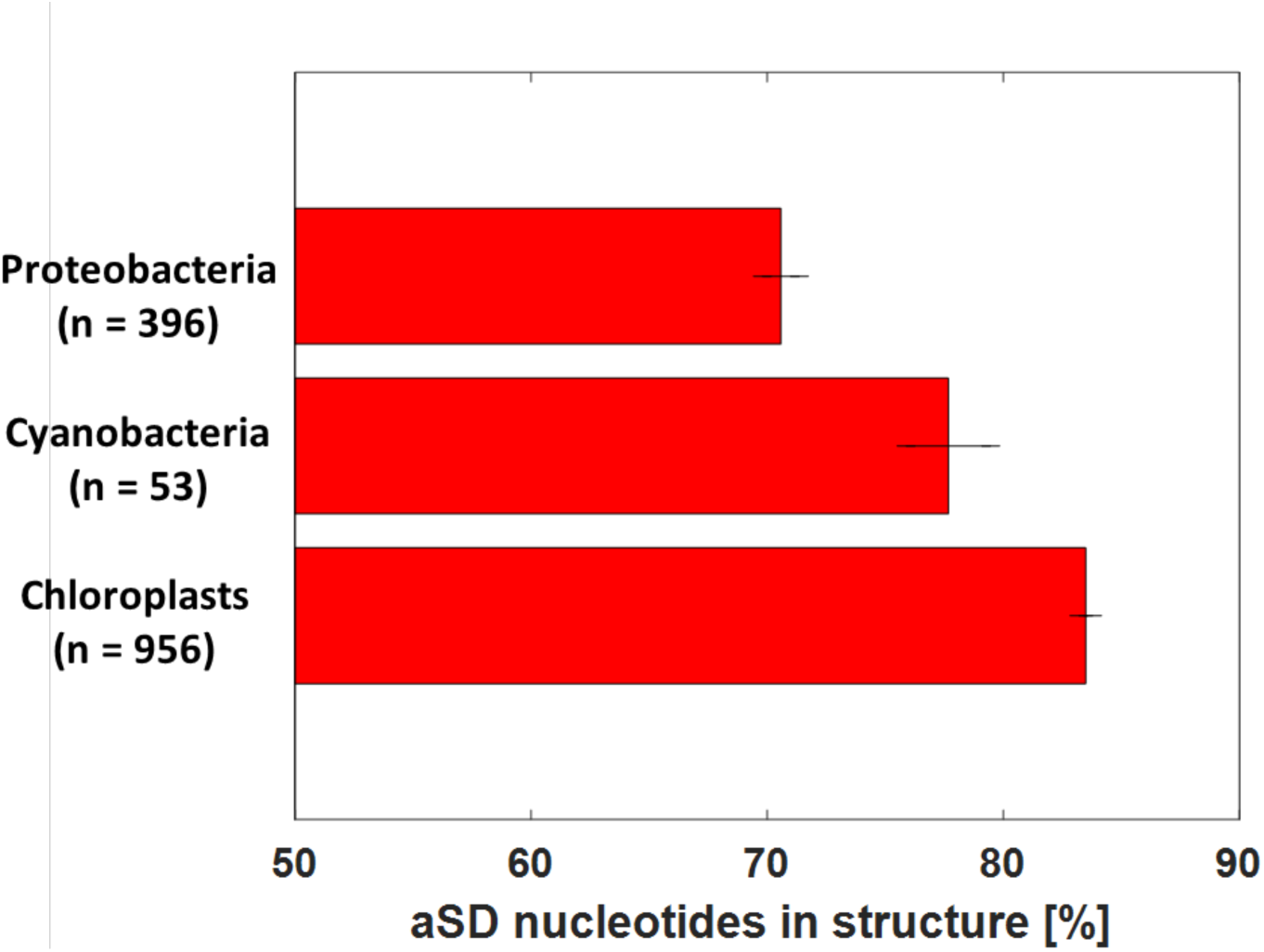
Counting nucleotides in structure while assuming a 6-base consensus sequence. This is the same analysis as in Fig. 2*C*, where only the Shine-Dalgarno consensus sequence chosen was AGGAGG (instead of AGGAGGU). The main conclusion from this test – that chloroplast and cyanobacteria anti-Shine-Dalgarno sequences are bound by more base-pair interactions than their proteobacterial counterparts – remains similar, regardless of the consensus sequence definition.

